# Cleavable crosslinkers redefined by novel MS3-trigger algorithm

**DOI:** 10.1101/2023.01.26.525676

**Authors:** Lars Kolbowski, Lutz Fischer, Juri Rappsilber

## Abstract

Crosslinking MS is currently transitioning from a routine tool in structural biology to enabling structural systems biology. MS-cleavable crosslinkers could substantially reduce the associated search space expansion by allowing an MS3-based approach for identifying crosslinked peptides. However, MS2-based approaches currently outperform approaches utilising MS3. We show here that MS3-trigger sensitivity and specificity were hampered algorithmically. Our four-step MS3-trigger algorithm greatly outperformed currently employed methods and comes close to reaching the theoretical limit.

## Main

Crosslinking mass spectrometry (MS) is a discovery tool of hitherto hidden aspects of biology^1^, from placing protein sequence in unassigned densities of cryoEM and cryoET data^2^ to capturing weak interactions in whole cells that are lost upon lysis^3^. Among the plethora of crosslinking MS workflows^4–6^, the use of MS-cleavable crosslinkers stands out^7–12^. Their cleavage reverses the crosslink in the mass spectrometer, such that the two peptides can be analysed individually. While this can also be achieved computationally during data analysis^13^ doing so in the mass spectrometer improves peptide fragmentation^14^. Accordingly, cleavable crosslinkers improve the number of reliably identifiable crosslinks, especially in complex samples^14,15^. In principle, the unlinked peptides could also be selected for separate fragmentation in MS3^16^. MS3 would tremendously simplify data analysis, however, most investigations focus on MS2 data^17,18^. The additional acquisition time cost due to MS3 spectra acquisition is one of the downsides of MS3-based approaches^14,19^ that would need to be counterbalanced by clear advantages. However, MS3 approaches based on MS-cleavable crosslinkers are currently limited by two technical challenges^14^. While the signature doublets of the two peptides can routinely be detected in the spectra of crosslinked peptides (81%), only 41% of the individual peptides are selected for MS3 (low sensitivity). In addition, much time is wasted on MS3 of not crosslinked peptides that are abundantly present in the analysed samples (low specificity). We here find the culprit in the current algorithms used for MS3 decision-making and present a novel algorithm for MS3-based acquisition strategies.

We designed a four-step procedure that was highly likely to improve results based on our previous analysis^20^. First, isotopic envelopes in the spectrum are detected, deconvoluted to monoisotopic peaks with summed intensities, and assigned charge states (**Fig. 1a**). Second, for each peak with a defined charge state, its theoretical doublet partner peak is calculated by the addition of the doublet delta mass divided by the peak’s charge state. These theoretical doublet peaks are then matched against peaks with matching charge states, considering a user-defined relative *m/z* tolerance (**Fig. 1b**). Third, a rank-based cut-off is applied, disregarding all doublets with a rank smaller than 20, to reduce noise matching (**Fig. 1c**). Fourth, we implemented a ‘second peptide mass’ filter, throwing out doublets that leave less than 500 Da for the second peptide, resulting in the final matched doublet list (**Fig 1d**). If the remaining mass is less than 500 Da, no MS3 should be acquired as the doublet stems either from a linear peptide with a crosslinker modification, is a false positive random match, or the second peptide is too small to be reliably identifiable (**Fig. S1**). We chose 5 ppm as the mass tolerance for matching the doublet delta mass, according to the data accuracy and balancing specificity and selectivity (**Fig. S2**).

**Figure 1.**
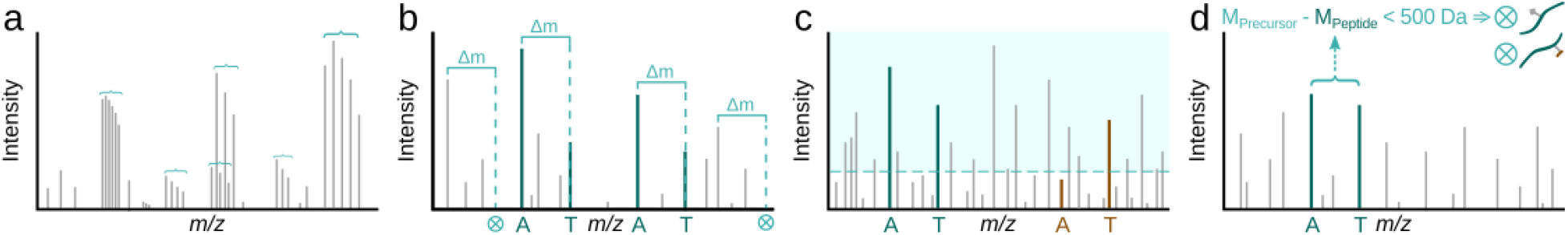
Flowchart of the improved MS3-trigger algorithm. (a) Deconvolution of isotope clusters and charge determination. (b) Doublet matching by Δm and charge state. (c) Intensity rank filter. (d) Remaining second peptide mass filter.

Each matched doublet from the final output of our algorithm would have triggered the acquisition of an MS3 spectrum. We evaluated this against the actual triggered MS3 from the acquisition with regard to specificity and sensitivity. In the best case of sensitivity, we would expect to trigger an MS3 for a doublet of every peptide that we know from annotating the identified CSMs. This gives a theoretically possible maximum of 85% (**Fig. 2a**, Synaptosome) and 76% (**Fig. 2d**, Ribosome) of all CSMs. The correct trigger rate improved on both peptide doublets from 58% to 79% and from 37% to 60% of CSMs for the Synaptosome and Ribosome dataset, respectively, thus closing the gap to the theoretical maximum (**Fig. 2b,e**). Furthermore, our doublet selection algorithm vastly improved the specificity in both datasets (**Fig. 2c,f**). We could almost completely eliminate MS3 triggers in (linear) PSMs (−92%). On average, only 0.11 and 0.09 MS3 spectra were triggered in the Synaptosome and Ribosome datasets for these not crosslinked peptides. In addition, MS3 triggers for crosslinker-modified PSMs were also reduced substantially by approximately 75% on average.

**Figure 2.**
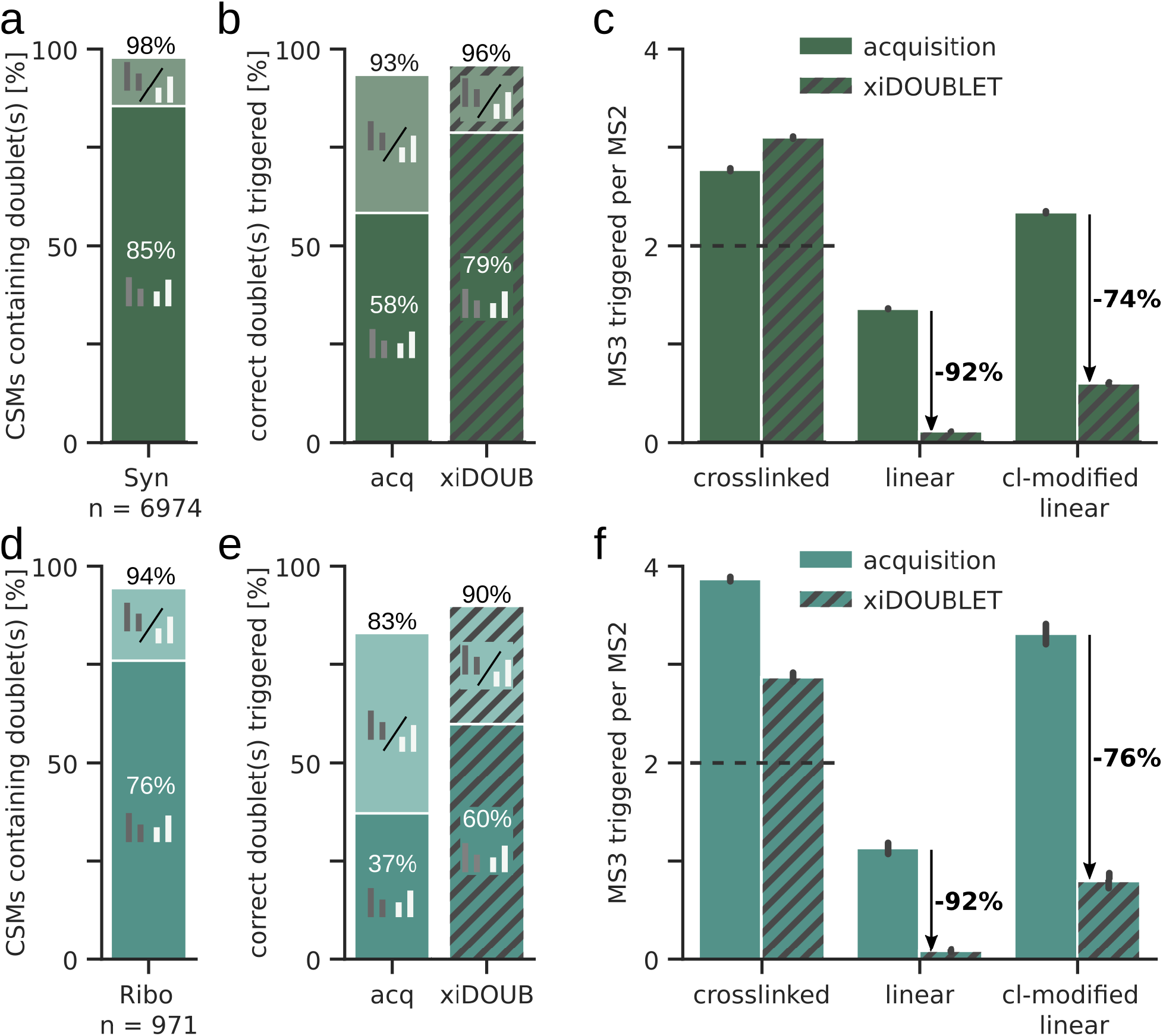
Sensitivity and specificity of MS3 triggering. (a, d) Proportion of identified CSMs that contain one (lighter colours) or both (darker colours) peptide doublets in each dataset (5% CSM-level FDR). (b, e) Proportion of correctly triggered MS3 scans acquisition data compared to the results from using in-silico triggering using the xiDOUBLET algorithm. (c, f) Number of triggered MS3 scans per MS2 scan, comparing DSSO acquisition results to the xiDOUBLET algorithm. Ideally, two MS3 scans are triggered for a crosslink (one for each of the two peptides, dotted line) and none for linear and modified linear peptides. Error bars show the 0.95 confidence interval. Panels (a - c) represent the Synaptosome, panels (d - f) the Ribosome data.

Gains in sensitivity and via time savings during acquisition also gains in selectivity greatly affect the analytical depth. We demonstrate here that sensitivity and specificity can be improved considerably with no change in the experimental side of established CLMS protocols. Our four-step procedure vastly outperforms the doublet selection algorithm currently employed on Orbitrap mass spectrometers. The closed source code of the instrument control software prevents us from implementing our procedure, making a case for open code in the interest of scientific progress and extending an urgent call on the vendor to implement the improved procedure. Our work shows, nonetheless, that MS3 approaches suffer from algorithmic restrictions that can be overcome to help unfold the full potential of MS-cleavable crosslinkers for structural proteomics.

## Acknowledgements

This research was funded by the Deutsche Forschungsgemeinschaft (DFG, German Research Foundation) under Germany’s Excellence Strategy – EXC 2008 – 390540038 – UniSysCat and project 426290502 and, in part, by the Wellcome Trust [Grant number 203149]. For the purpose of open access, the authors have applied a CC BY public copyright license to any Author Accepted Manuscript version arising from this submission.

## Supporting Information

**Figure S1.**
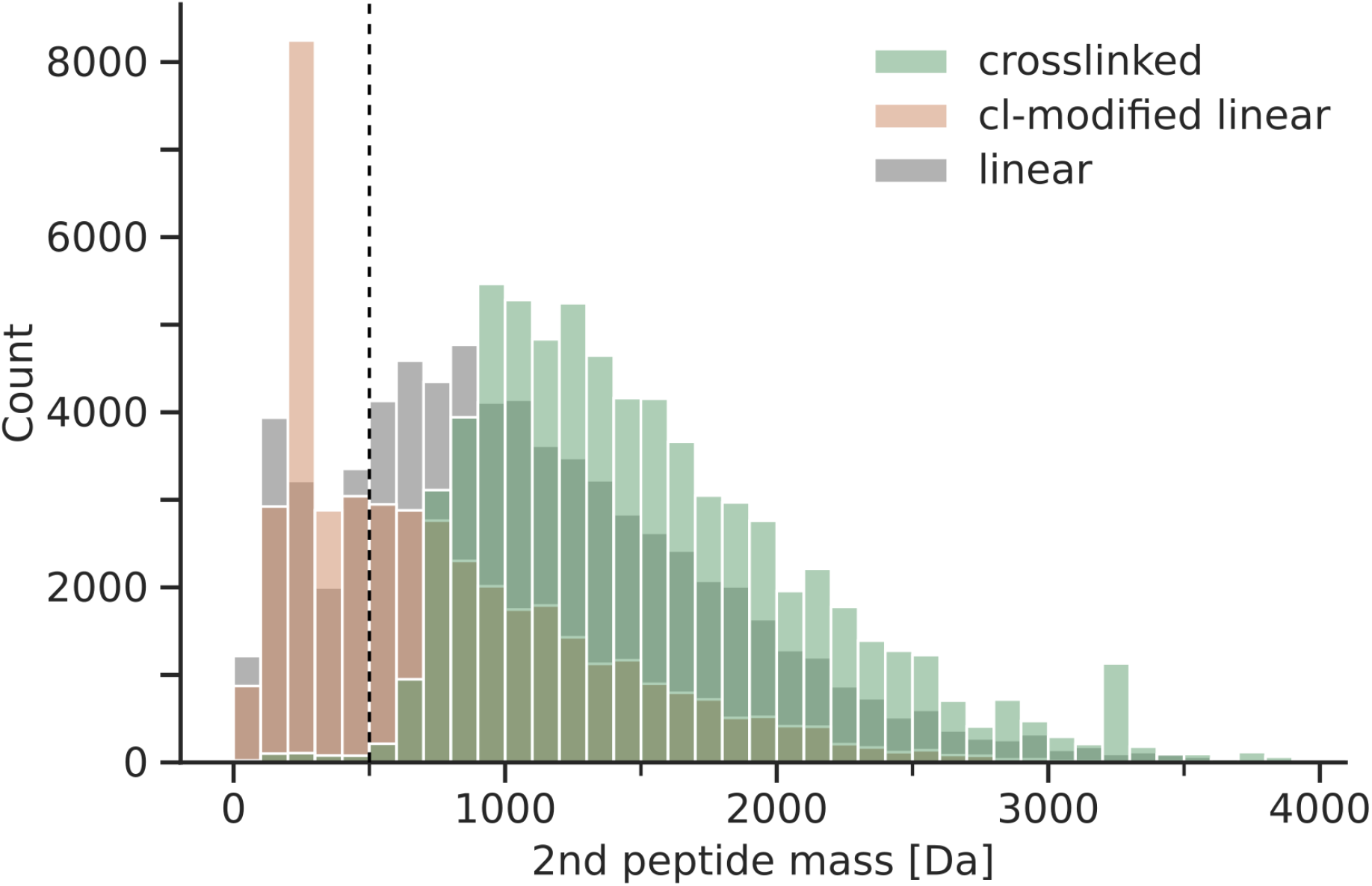
Histogram of remaining second peptide masses for all doublets (filtered to top 20 peaks and capped to the max. 4 highest ranking doublets per spectrum) detected in identified CSMs (green) and linear PSMs with crosslinker modification (orange) and without (grey) from both datasets. The dashed line shows the chosen cut-off.

**Figure S2.**
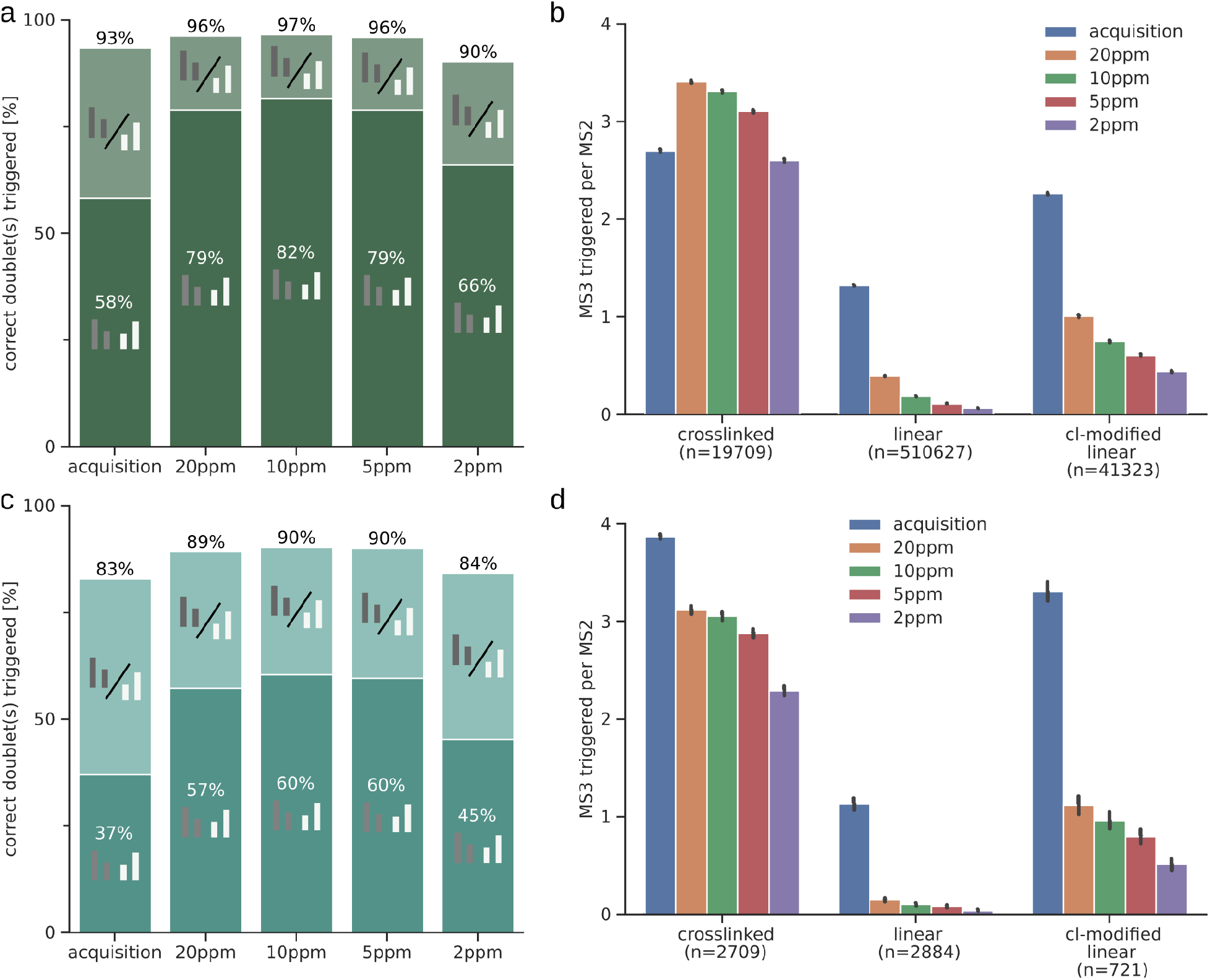
Sensitivity and specificity of MS3 triggering using different mass tolerances. (a, c) Proportion of correctly triggered MS3 scans acquisition data compared to the results from using in-silico triggering using the xiDOUBLET algorithm. (b, d) Number of triggered MS3 scans per MS2 scan, comparing DSSO acquisition results to the xiDOUBLET algorithm. Error bars show the 0.95 confidence interval. Panels (a - b) represent the Synaptosome, panels (c - d) the Ribosome data.

## Methods

### Data Analysis

We reanalysed the publicly available CID MS3-based data from the Ribosome (PXD011861) and Synaptosome (PXD010317 & PXD015160) datasets. The datasets were analysed as previously described^14^. A 5% CSM level FDR, with sequence-consecutive and minimum peptide length (5 amino acids) filters, was applied. The CID spectra were annotated using pyXiAnnotator according to the CSM identification with b-, and y-ion series and the cleavable crosslinker stub fragments A, S, and T using a 15 ppm fragment mass tolerance. For the doublet rank evaluation, the “deisotoped max rank” column from pyXiAnnotator output was used which determines the rank of the annotated isotope cluster by comparing the maximum intensity peak of each isotope cluster. Doublet rank was then assigned by the higher of the two doublet peak ranks. To evaluate if the correct peaks were triggered for MS3, the MS3 precursor *m/z* was extracted from the scan header of MS3 spectra associated with the unique CSMs passing FDR (as described above) and compared with the corresponding MS2-CID annotation result. If the MS3 precursor matched a crosslinked peptide stub fragment within 20 ppm error tolerance, it was assigned as correctly triggered. For the evaluation of the MS3 trigger specificity, the number of MS3 scans associated with non-unique CSMs and linear PSMs (with and without hydrolyzed or amidated crosslinker modifications) passing the FDR threshold was used.

The settings for the xiDOUBLET doublet detection algorithm used here were: ms2_tol of 5 ppm tolerance; crosslinker DSSO; stubs A & T; rank_cutoff of 20; cap of 4; second_peptide_mass_filter 500. The algorithm is written in python and is open source and freely available on https://github.com/Rappsilber-Laboratory/xiDOUBLET.

